# Neural Mechanisms Linking Global Maps to First-Person Perspectives

**DOI:** 10.1101/2025.03.03.641106

**Authors:** Hin Wai Lui, Elizabeth R. Chrastil, Douglas A. Nitz, Jeffrey L. Krichmar

## Abstract

Humans and many animals possess the remarkable ability to navigate environments by seamlessly switching between first-person perspectives (FPP) and global map perspectives (GMP). However, the neural mechanisms that underlie this transformation remain poorly understood. In this study, we developed a variational autoencoder (VAE) model, enhanced with recurrent neural networks (RNNs), to investigate the computational principles behind perspective transformations. Our results reveal that temporal sequence modeling is crucial for maintaining spatial continuity and improving transformation accuracy when switching between FPPs and GMPs. The model’s latent variables capture many representational forms seen in the distributed cognitive maps of the mammalian brain, such as head direction cells, place cells, corner cells, and border cells, but notably not grid cells, suggesting that perspective transformation engages multiple brain regions beyond the hippocampus and entorhinal cortex. Furthermore, our findings demonstrate that landmark encoding, particularly proximal environmental cues such as boundaries and objects, play a critical role in enabling successful perspective shifts, whereas distal cues are less influential. These insights on perspective linking provide a new computational framework for understanding spatial cognition and offer valuable directions for future animal and human studies, highlighting the significance of temporal sequences, distributed representations, and proximal cues in navigating complex environments.

**Significance Statement:** Understanding how the brain transforms between different spatial perspectives is crucial for advancing our knowledge of spatial cognition and navigation. This study presents a novel computational approach that bridges the gap between neural recordings and behavior, offering insights into the underlying mechanisms of perspective transformation. Our findings suggest how the brain integrates temporal sequences, distributed representations, and environmental cues to maintain a coherent sense of space. By demonstrating the importance of proximal cues and temporal context, our computational model provides testable predictions for future neurophysiological studies in humans and animals.

---

Humans and other animals have the ability to navigate familiar environments by observing their surroundings in a first-person perspective (FPP). That is, we know where we are by just looking at our surroundings, and we are able to plan a route and navigate to our end goals. This navigation ability sometimes requires the navigator to use a different perspective in a broader context, as if one is looking at oneself from a bird’s-eye, top-down perspective, which we call the global map perspective (GMP).

Indeed, evidence suggests that humans and many animals have this ability to transform locations in the FPP to the same locations in the GMP (1–4). On the other hand, evidence suggests that the reverse is also true; that humans and other animals are able to seamlessly execute transformations from GMP to FPP. For example, when a squirrel in a tree sees an acorn on the ground below (GMP), it moves to ground level, and then navigates to pick it up using the FPP. In particular, bats appear to have the ability to translate a GMP while flying above the landscape to an FPP when navigating on the ground or foraging (4–7).

Numerous studies have explored the neural mechanisms of the brain that perform perspective transformation. Given the spatial and directional tuning correlates of their constituent neuron populations, it is likely that the entorhinal cortex (EC), retrosplenial cortex (RSC), subiculum (SUB), posterior parietal cortex (PPC), and hippocampus (HPC) play significant roles in linking locations and orientations relative to one perspective to the same locations and orientations relative to another perspective (1, 8–13). In rodent navigation studies with ensemble neuron recordings of individual cells, many different cell types are found to be highly tuned to different spatial and environmental features that are required for successful navigation. For example, place cells in the hippocampus are active when the animal is in specific locations within an environment (14–16), head-direction cells in the RSC, medial entorhinal cortex (MEC), and subiculum are highly tuned to specific head directions (17, 18), grid cells in the entorhinal cortex fire maximally in a spatially repeating pattern when the animal freely navigates the environment (19). Further, boundary vector cells in the subiculum (20) and object vector cells in the MEC (21) fire maximally when the animal is at a specific distance and orientation to an environment boundary or object.

It is important to note that both first-person perspective (FPP) and global map perspective (GMP) involve allocentric representations. The transformation between these perspectives is not simply a shift between egocentric and allocentric frames, but rather a complex interaction of multiple allocentric representations. In both cases, localization within these allocentric frames is achieved through learning associated egocentric representations while an animal navigates its environment.

Several theories have emerged to explain how these cell types might facilitate the transformation between perspectives, including the integration of multiple potential allocentric representations - a process we call *perspective alignment* (1, 9, 11, 13). However, the exact computational mechanisms that mediate such perspective alignment remain to be defined and tested. Additionally, the forms of spatial and directional tuning that best support such computations have yet to be identified. Our work aims to address these open questions by investigating the computational principles underlying perspective transformations, without presupposing the existence of specific cell types or representations.

In our previous work (22), which made no assumption on the underlying mechanism of perspective transformation, we demonstrated successful performance of perspective transformations between FPPs and GMPs. We also used variational autoencoders (VAEs) (23) to perform perspective transformation between GMPs and FPPs, and vice versa, on paired images obtained when a robot is freely navigating a simulated environment. The VAE reconstructed perspective-transformed images with high detail and fidelity, using two separate VAEs for each direction. By analyzing the tuning curves of the latent variables of the VAEs, we found the emergence of features resembling place cells and head-direction cells found in rodent neuron recording experiments, even though these features were not part of the VAE itself.

One limitation of that work was that the perspective transform network was trained on static pairs of images from both perspectives, without any temporal or sequential components to the model. However, this approach is biologically unrealistic, because humans and other animals navigate and update their mental location in a continuous manner. Another limitation was that only place cells and head direction cells were investigated without considering other prominent cell types, such as grid cells, border cells, corner cells, and object vector cells. Finally, the visual cues used by the model to perform these transformations were not investigated. For example, it was unclear whether the model used more proximal or distal cues to perform perspective transformations.

In the present work, we utilize recurrent neural networks (RNNs) (24) as the latent variables of the VAE. Each RNN processes a sequence of images from one perspective and generates the associated sequence for another perspective. RNNs maintain an internal state that evolves with changing inputs, allowing for sequential output generation. This approach enables a more realistic modeling of the perspective transformation process, conceptualized as a sequence of input and output images. We hypothesized that this sequential processing would better capture the temporal aspects of spatial navigation and lead to more biologically plausible representations.

Importantly, we did not explicitly incorporate any specific cell types (such as place cells or grid cells) into our model architecture. Instead, we aimed to observe whether such representations would emerge naturally from the task of perspective transformation. To this end, we added an attention block, which allows the model to selectively attend to different parts of the input image, to identify the cues and stimuli critical to these transformations. We then analyzed the latent variables for properties resembling various cell types, including grid cells, boundary vector cells, and object vector cells. This analysis provides a richer understanding of how complex spatial representations might arise and interact during viewpoint transformations.

Finally, we conducted causal perturbation and ablation studies to test our hypothesis that proximal cues would be more important than distal cues for perspective transformation. These studies allowed us to systematically investigate the relative importance of different environmental features in our model’s performance.

Through this approach, we aimed to shed light on the computational principles that might underlie the brain’s ability to perform perspective transformations, without presupposing the existence of specific cell types or representations.

## Results

### Robotic Simulation and Model Architecture

Similar to (22), we used the Webot robot simulation environment (25, 26) to collect data for the training of the VAE. A Khepera robot with a head-mounted camera moved freely within an arena with three colored cylinders (Fig. 1a). Every 20 frames, we collected a pair of images from both the FPP camera and the GMP map. We collected 10,000 pairs of images in total, with 8,000 images used for training and validation of the VAE, and 2,000 images for testing. All latent variable analysis were from these 2,000 test images. We repeated each model with different configurations 5 times with 5 different random seeds. Differing from (22), we used colored walls with distinct colors for each corner to provide additional location-specific visual cues, as well as using a non-checkerboard floor pattern to prevent a grid-like response from the latent variables of the VAE due to simply the floor pattern. Because the robot tended to spend more time near walls and corners, we introduced random location targets in the arena for the robot to visit. This is similar to the common practice of randomly distributing food rewards in rodent experiments to promote exploration.

**Fig. 1.**
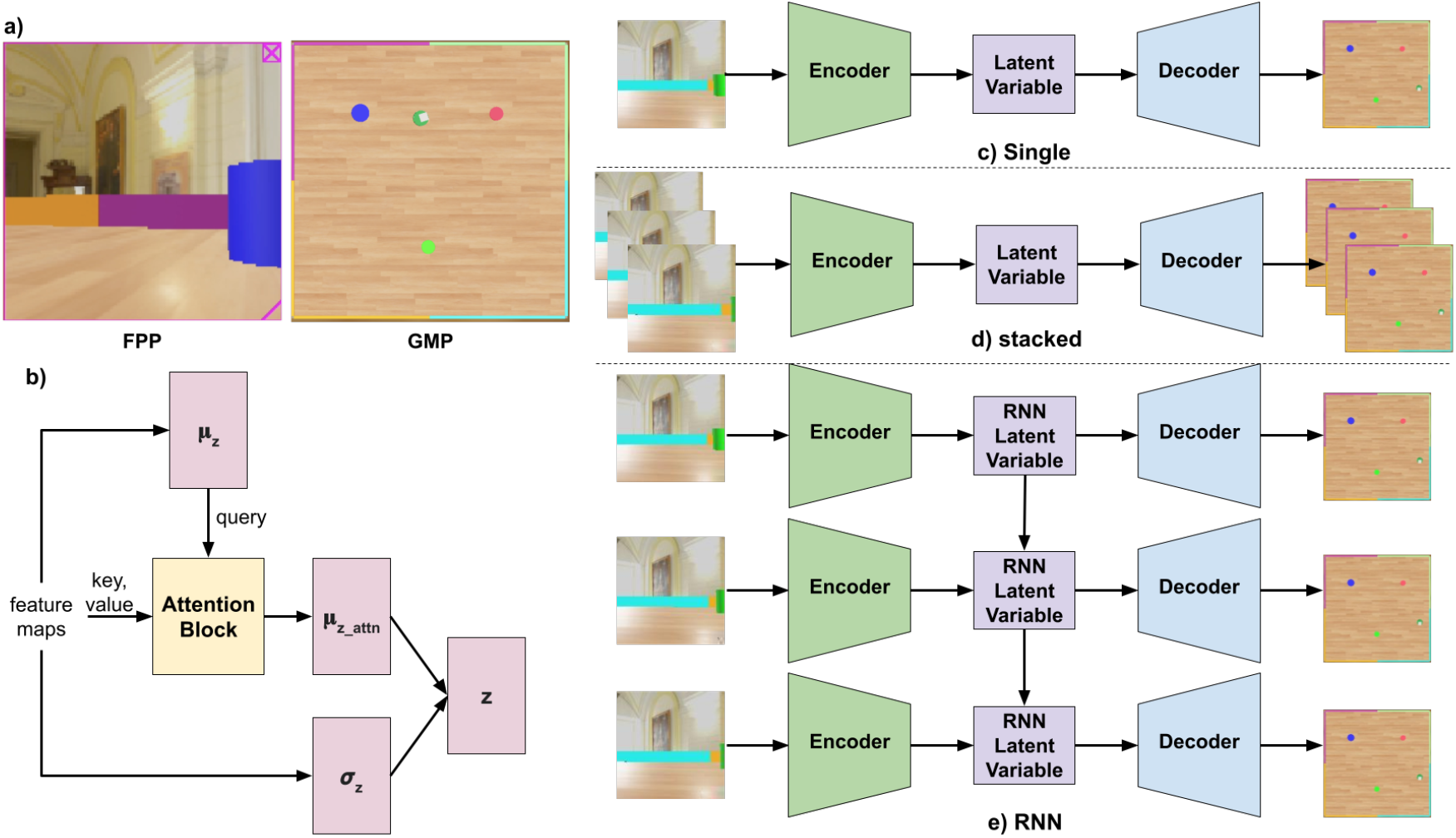
Experimental setup and model architectures for perspective transformation. (a) Webot simulation environment showing first-person perspective (FPP, left) and global map perspective (GMP, right) of a Khepera robot freely exploring a 1×1 meter arena with colored walls and cylinders. (b) Convolutional Block Attention Module (CBAM) for latent variables, where *µ*_*z*_ and *σ*_*z*_ are the mean and variance of a Gaussian distribution generating latent variable *z*. CBAM allows *µ*_*z*_ to attend to specific environmental stimuli, resulting in *µ*_*z attn*_. (c-e) Schematics of three VAE architectures: (c) “Single” with individual image input and output, (d) “Stacked” with a sequence of images as input and output, allowing simultaneous processing of multiple frames, (e) “RNN” with recurrent neural network as VAE latent variable, processing and generating image sequences. All architectures perform perspective transformation between FPP and GMP in both directions as separate models. The number of frames in the input sequence varies across experiments, allowing for investigation of temporal effects on spatial representation.

We compared three different model architectures (Fig. 1). Each was constructed to allow for an “attention block” to examine latent variable responses to environmental stimuli (Fig. 1b). In Fig. 1c, the *Single* VAE reconstructed one perspective (e.g., GMP) from the other (e.g., FPP) one image at a time without any temporal information. This is the same model used in (22). In Fig. 1d, the *Stacked* VAE reconstructed a sequence of images of one perspective, from the sequence of images from another perspective. In Fig. 1e, the *RNN* VAE reconstructed one perspective from the other using recurrently connected latent variables. In this model, only the latent variables were recurrent, while the encoder and decoder of the VAE were non-recurrent. It was trained end-to-end with backpropagation through time. For each time step in the sequence, it took the input image from one perspective and generated an image for another perspective. Between each time step, the hidden states of the RNN were recurrent, meaning that they were carried forward in time to the next pair of images.

The difference between the stacked and RNN model was that the former uses information from all the input images to reconstruct all the output images at the same time, whereas the latter reconstructs images one step at a time, taking account of only the current input image and the previous state of the latent variable. We kept the same FPP ↔ GMP correspondence of images between the two perspectives for all models. This ensured the dimensionality between the inputs and the outputs was the same across all models, ensuring a fair comparison. Fig. 2a shows examples of the source image to the VAE, the target output, and the reconstructed images from the models for both transformation directions.

**Fig. 2.**
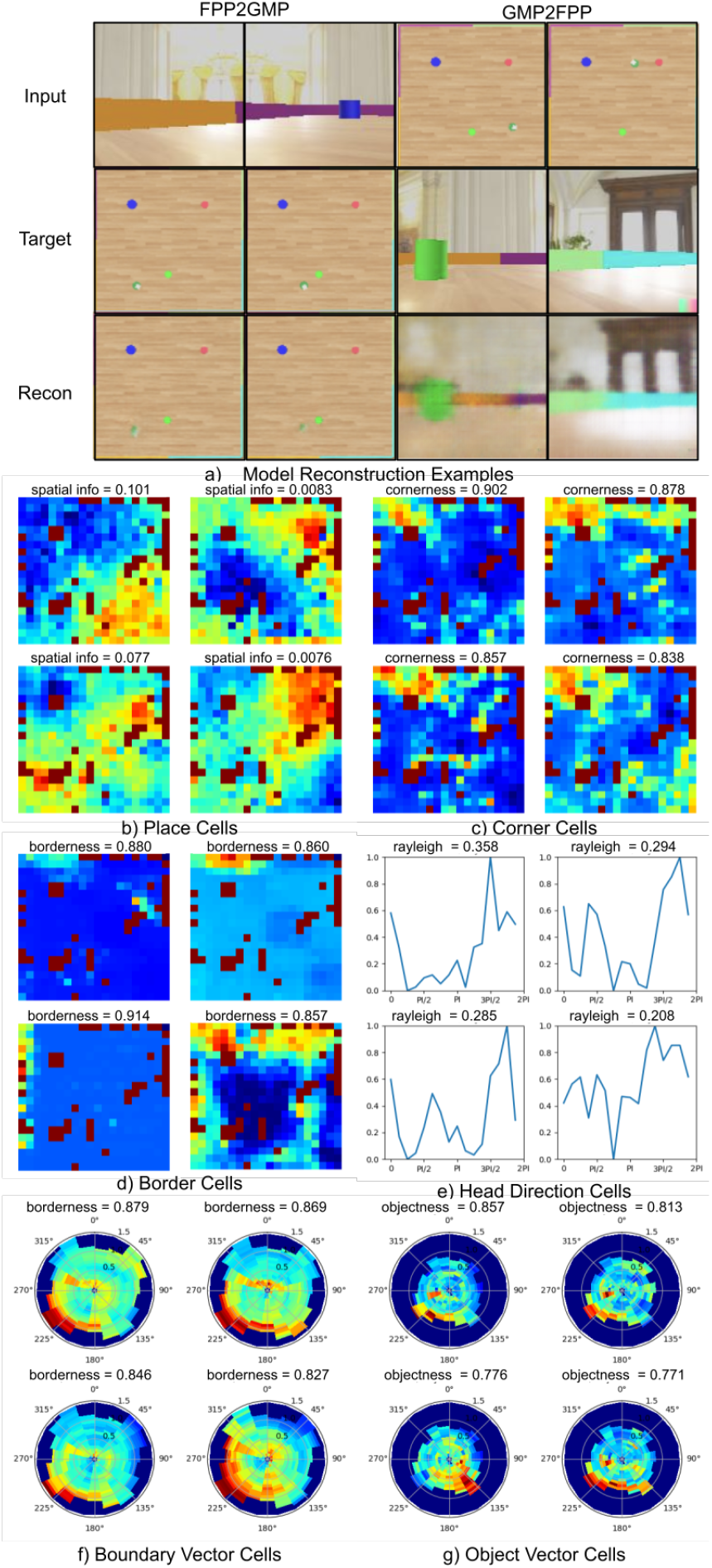
Emergence of biologically-inspired spatial representations in the RNN-VAE model. (a) Example of images reconstructed by the RNN-VAE, showing source, target, and model output for both FPP to GMP and GMP to FPP transformations. (b-g) Latent variables of the VAE exhibiting response properties similar to various spatial cell types found in rodent experiments, ranked by their corresponding spatial metrics (left to right, top to bottom): (b) place cells, (c) border cells, (d) corner cells, (e) head direction cells, (f) boundary vector cells, and (g) object vector cells. These cells emerged without explicit programming, suggesting the model captures fundamental computational principles underlying spatial cognition.

In addition to the RNN-VAE architecture, we incorporated an attention mechanism into our model to enhance its ability to focus on relevant features during perspective transformation. This mechanism, implemented as a Convolutional Block Attention Module (CBAM, Fig. 1b)(27), allows the model to selectively attend to different parts of the input image. The CBAM takes the mean (*µ*_*z*_) of the variational Gaussian distribution of the VAE as a query, and the image feature maps from the encoder as keys and values. This setup enables each latent variable to pay specific attention to important features in the spatial feature maps. The attention mechanism operates by computing a weighted sum of the input features, where the weights are determined by the relevance of each feature to the current transformation task. This process mimics the brain’s ability to selectively focus on certain aspects of sensory input while ignoring others. By incorporating this attention mechanism, we aimed to provide our model with a more flexible and dynamic way of processing spatial information, potentially leading to more robust and interpretable perspective transformations. Furthermore, analyzing the attention patterns allows us to gain insights into which environmental features the model deems most important for successful perspective shifting, offering a window into the model’s decision-making process.

### Latent variable analysis

For the latent variables of the VAEs, we analyzed their tuning curves and responses to place, head-direction, and object-orientation, and found the emergence of key spatial navigational and spatial cell types also found in rodent experiments. Fig. 2(b)-(g) shows typical examples of latent variables from the RNN-VAE that resemble place cells, border cells, corner cells, head direction cells, boundary vector cells, and object vector cells.

We used a set of standard metrics commonly used within the neuroscience community to test how similar these latent variables were to the cell types. To quantify spatial tuning of place cells (Fig. 2b), we used spatial information and spatial coherence. Spatial information (16) measured the extent to which activity rates were high across a small subset of locations and were low or nonexistent across the remainder of an environment. Coherence (15) measured the extent to which high activity rates clustered in a single location.

For border cells Fig. 2d), we used the borderness score, which measured the peak average response at any of the 4 walls, adjusted by the average firing distance of the entire map to the wall. Lever et al. (28) provided extensive data characterizing these cells, which they termed boundary vector cells (BVCs). They demonstrated that these cells fire at a specific distance and direction from environmental boundaries, maintaining their firing fields across different environments. Lever’s work showed that BVCs are found in the subiculum and are distinct from hippocampal place cells, responding primarily to the presence of boundaries rather than to specific places. In a follow-up study, Lever et al. (29) further elucidated the properties of BVCs, showing their rapid formation in novel environments and their stability across different behavioral states. In our model, the emergence of border cell-like activity in the latent variables suggests that boundary information is crucial for the task of perspective transformation between first-person and global map views. This aligns with the biological findings that border cells play a significant role in spatial representation and navigation.

For corner cells (Fig. 2c) and object cells (Fig. 2g), we used the same method as the borderness score to compute the cornerness score and the objectness score, except the pixels of interest changed from those immediately adjacent to the walls, to those pixels adjacent to the 4 corners and the 3 cylinder objects.

For head direction cells (Fig. 2e), we used the head direction score (30) and Rayleigh score (31), which measured how the head direction tuning curve differed from a uniform distribution around the circle.

We also used gridness score (32) to search for grid cells. Gridness measures the ratio of peak to trough response of the spatial auto-correlation map rotated at different angles. However, results show that there were no latent variables with high enough grid score to be classified as grid cells.

The latent variables of the VAE acted as an information bottleneck of the encoder-decoder generative model. Effectively, they performed a non-linear dimensionality reduction of the input images, similar to principal component analysis, by compressing the image data with 128 *×* 128 pixels to only 50 latent variables. Therefore, the latent variables represented the high-level information within the input images that were representative of the task of generating the images for another perspective. The fact that the VAE models were able to generate the images of the corresponding perspective with high detail, fidelity, and accuracy meant that the models were successful at finding the key factors within the environment that were indicative of the position of the robot in space. The emergence of the cell types found in Fig. 2 suggests that, from an information processing perspective, head direction, distance, and orientation to environment boundaries and key objects were all important factors for successful localization in space.

### Comparing Sequential Architectures

We evaluated and compared the relative performance and the quality of the latent responses between the three different neural network architectures: 1) single, 2) stacked, and 3) RNN. In addition to the three architectures, we include the results of an untrained RNN model with random weight initialization (red line in Fig.3), to ensure the latent variable responses obtained for the trained models were direct results of learning the spatial structural basis required for perspective transformation. We compared the spatial information, coherence, and head direction scores. Additionally, we investigated the ability of the latent variables to capture the true position of the robot and how this ability was differed from each architecture. We trained a 2-layer multi-layer perceptron (MLP) model (33) to decode the position of the robot from its latent variables, and measured the distance deviation in meters, between the predicted and the ground truth position.

We found a greater spatial information for the RNN architecture than the single and the stacked architecture, as well as the randomly initialized untrained model, in both transformation directions (Fig. 3a). The stars in Fig. 3 indicate the significance level for the Wilcoxon rank-sum tests. The RNN had significantly higher spatial information than the other two architectures in the FPP to GMP direction and for the GMP2FPP direction compared to the stacked architecture. The architectures were not different from each other in spatial coherence in the FPP to GMP direction, but the RNN architecture had the higher extreme values in spatial coherence in the GMP to FPP direction (Fig. 3b). For head direction response, the RNN was better than all other architectures in both directions (Fig. 3c). The RNN’s latent variables were also able to better capture the true position of the robot than the other two models, with lower position deviations (Fig. 3d). Because the RNN proved to be the overall best architecture for modeling the perspective transform process, all subsequent analyses used the RNN model.

**Fig. 3.**
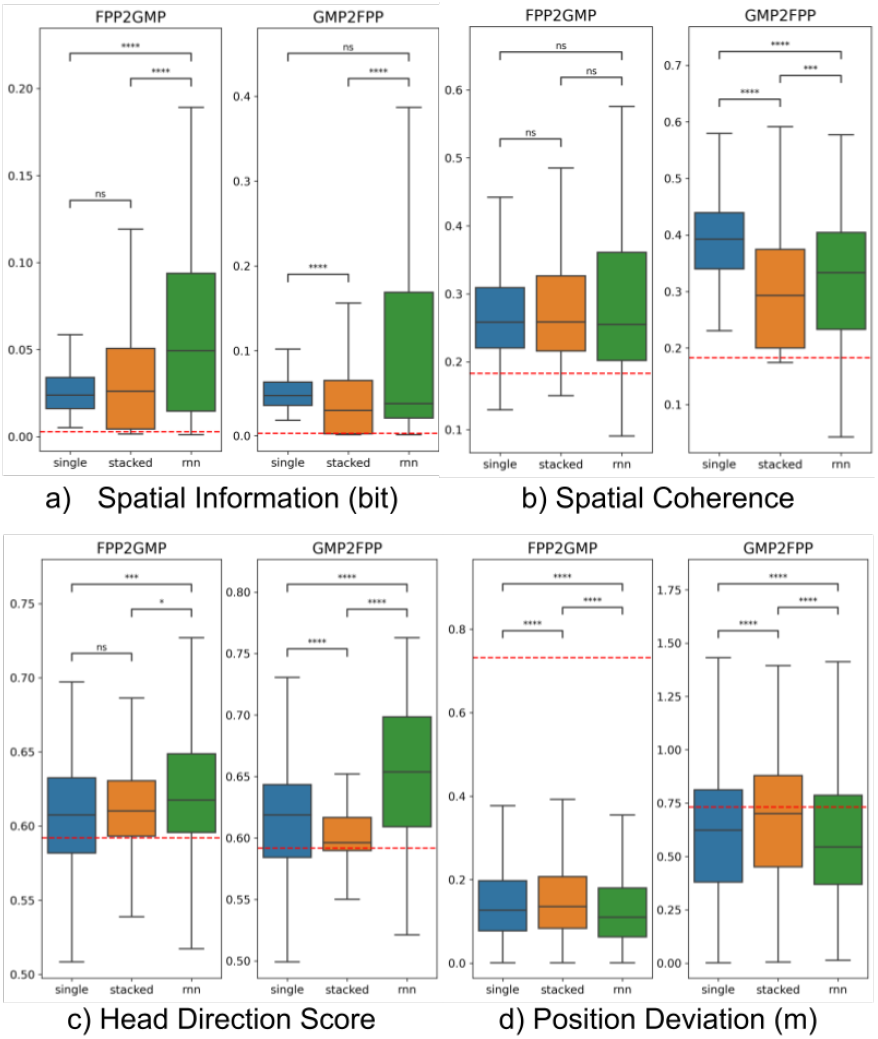
Performance comparison of different VAE sequential architectures on spatial metrics. Box plots show: (a) spatial information (bit), (b) spatial coherence, (c) head direction score, and (d) position decoding deviation (m) for Single, Stacked, and RNN architectures in both FPP to GMP and GMP to FPP directions. Red horizontal lines indicate mean performance of an untrained randomly initialized model. The RNN architecture consistently outperforms others, particularly in spatial information and head direction scores, demonstrating the importance of sequential processing in spatial representations. Statistical significance (Wilcoxon rank-sum test): * *p <* 0.05, ** *p <* 0.01, *** *p <* 0.005, **** *p <* 0.001, ns = not significant (34).

To further investigate the effect of incorporating sequential information into perspective transform modeling, we trained RNNs with varying sequence lengths, ranging from sequences of a single image (equivalent to the “single” architecture) to sequences of four images. For both perspective transformations, we observed a clear staircase effect on spatial information (Fig. 4a), where the spatial information increases as we move from RNNs trained with a single step to those trained with four steps in the sequence. This staircase effect is less pronounced for spatial coherence (Fig. 3b) and head direction score (Fig. 3c). Nevertheless, RNNs trained with longer sequences tend to display higher mean values or higher extreme values compared to those trained with just one time step. In terms of position decoding accuracy (Fig. 4d), RNNs trained with longer sequences demonstrate better position decoding than those trained with a single time step.

**Fig. 4.**
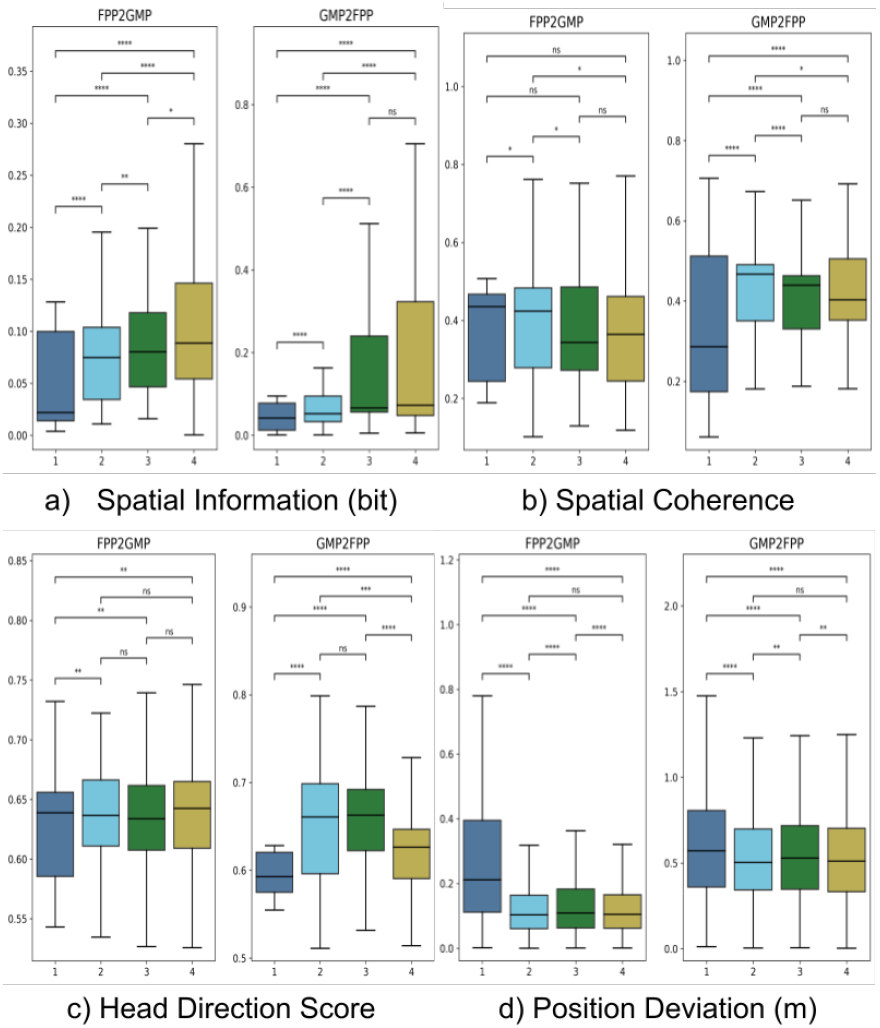
Effect of sequence length on RNN-VAE model performance. Box plots compare: (a) spatial information (bit), (b) spatial coherence, (c) head direction score, and (d) position decoding deviation (m) for RNN models trained with 1 to 4 time steps in both FPP to GMP and GMP to FPP directions. A clear improvement trend is observed as sequence length increases, particularly for spatial information and position decoding accuracy, underscoring the importance of temporal context in spatial representation. Statistical significance (Wilcoxon rank-sum test): * *p <* 0.05, ** *p <* 0.01, *** *p <* 0.005, **** *p <* 0.001, ns = not significant (34).

Overall, Figure 4 underscores the importance of incorporating sequential information into perspective transform modeling, enabling these models to perform perspective transformation tasks with greater biological resemblance.

### Comparison to Neural Correlates of Spatial Representations

Fig. 5 shows the histogram of the latent variables on different spatial metrics, with typical values for classifying particular cell types in rodent recordings highlighted in shaded green. Notably, we observed a consistent trend of stronger spatial representations in the Global Map Perspective (GMP) to First-Person Perspective (FPP) direction compared to the reverse.

**Fig. 5.**
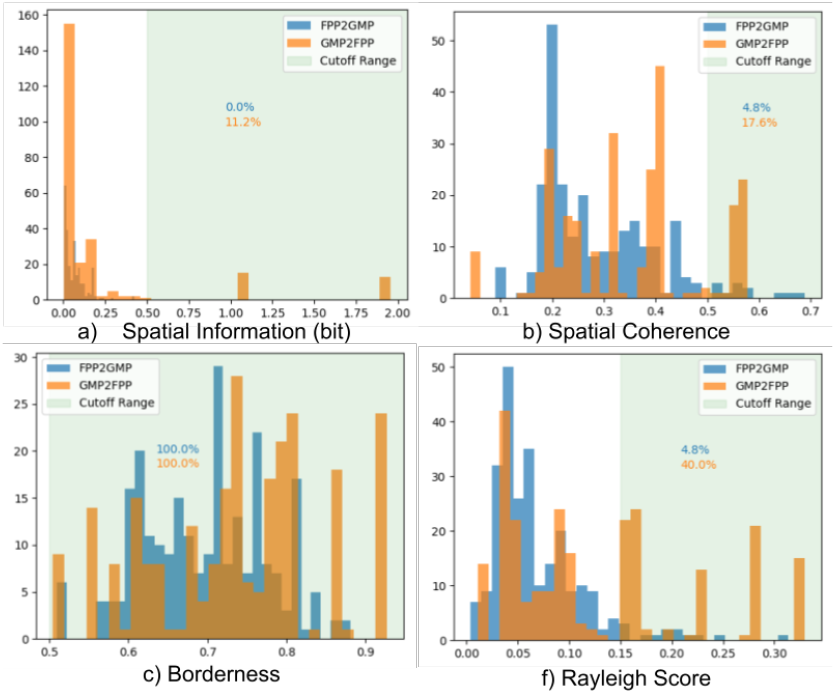
Comparison of RNN-VAE latent variable responses to rodent spatial cell criteria. Histograms show distribution of latent variables on different spatial metrics: (a) spatial information (bit), (b) spatial coherence, (c) borderness score, (d) head direction Rayleigh score. Shaded green areas indicate typical value ranges for classifying specific cell types in rodent experiments. Percentages of latent variables meeting these criteria are shown for both FPP to GMP and GMP to FPP directions. The model demonstrates a consistent trend of stronger spatial representations in the GMP to FPP direction.

For place cells, rodent hippocampal cells typically exhibit spatial information between 0.5-2.5 bits (16) and spatial coherence between 0.47-0.90. While 30% of hippocampus CA1 neurons are typically classified as place cells (35), our RNN model showed a directional asymmetry: in the GMP to FPP direction, 11.2% and 17.6% of the latent variables met the criteria for spatial information and coherence respectively, whereas these percentages were significantly lower in the FPP to GMP direction.

For border cells, typical values in rodent entorhinal cortex form a uniform distribution from −1 to 1, with a cutoff of 0.5 used to classify border cells, resulting in 11% meeting this criteria (30). Interestingly, our RNN model showed high borderness scores across both directions, with nearly all latent variables meeting this criteria.

Head direction cells in rodents typically have Rayleigh scores between 0 and 0.8, with a cutoff of 0.15 used to classify head direction cells, resulting in 44% of cells meeting this criteria in one study (32). Our RNN model closely matched this, with 40% of the cells meeting this criteria in the GMP to FPP direction. Again, we observed lower percentages in the FPP to GMP direction.

This consistent pattern of stronger spatial representations in the GMP to FPP direction suggests that our model may be capturing an important asymmetry in how the brain processes spatial information during perspective transformations. This directional preference aligns with the intuitive notion that it might be easier to generate a first-person view from a map-like representation than vice versa.

### Perturbation Study

To investigate whether distal or proximal features and cues were most important for spatial navigation and perspective transformation, we performed a series of perturbation studies. Fig. 6a shows the design of the perturbation studies. We rotated 3 layers of environmental features from proximal to distal and observed place field remapping. First, we rotated the three colored cylinders in the center 90 degrees counterclockwise (cylinder perturbation); next, we rotated the arena 90 degrees counterclockwise while keeping the cylinders the same as original (arena perturbation); finally, we rotated the distal background images 90 degrees counterclockwise while keeping everything else the same as the original (background perturbation). For background perturbation, due to technical constraints of the Webot robotic simulation environment not supporting the rotation of background images, we rotated the cylinder, the arena, and the coordinate system 90 degrees clockwise instead, making it equivalent to rotating the background image 90 degrees counterclockwise. For each perturbation, we performed a separate simulation and collected 2000 images from both perspectives and used the VAE-RNN models trained on the unperturbed version of the environment to test on the perturbed environments.

**Fig. 6.**
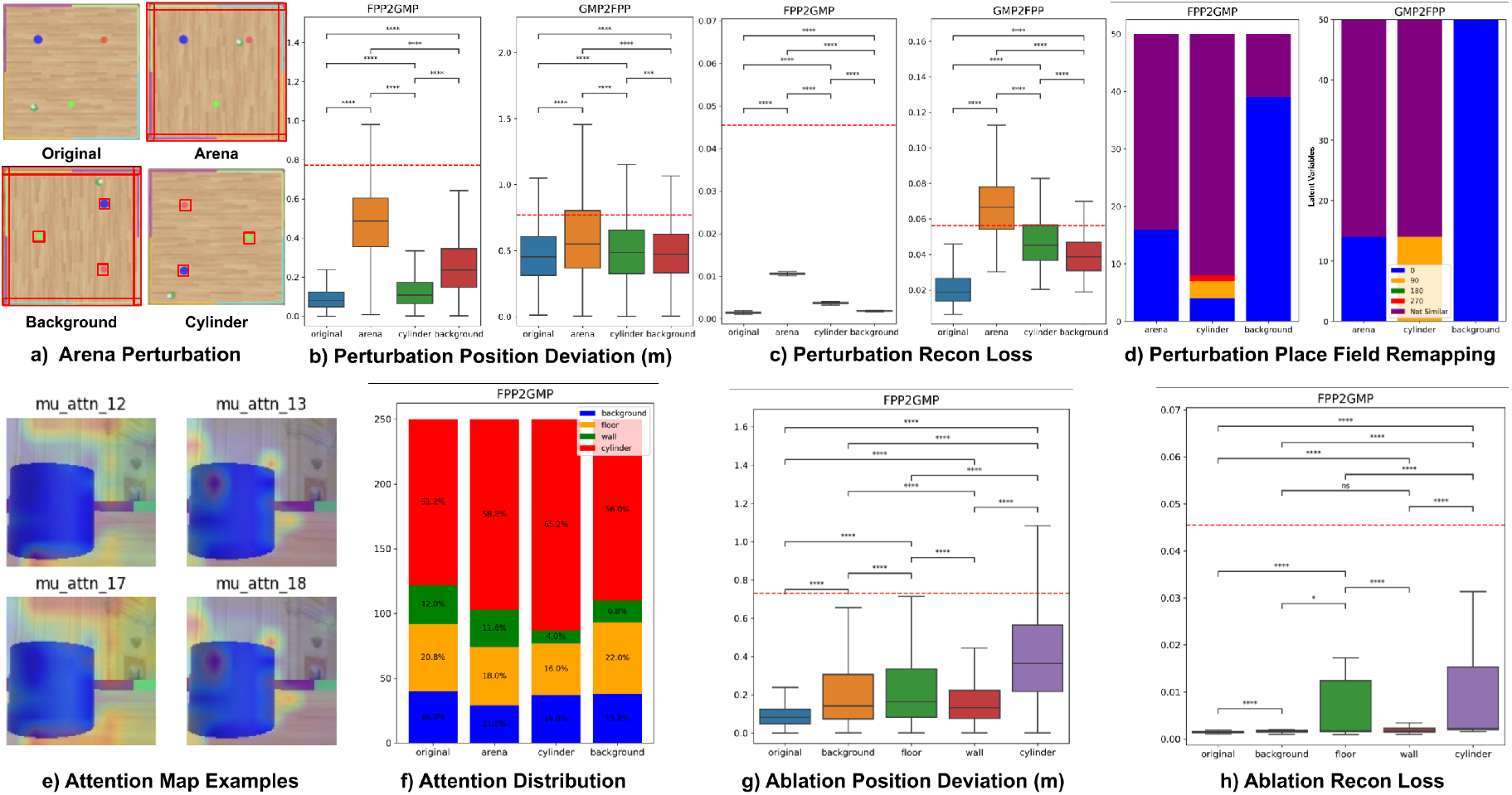
Perturbation and ablation studies revealing the importance of proximal cues in perspective transformation. (a) Schematic of perturbation experiments: arena, cylinder, or background rotated 90°counterclockwise (background implemented by rotating everything else clockwise, red box highlights perturbation). (b-d) Key performance metrics: (b) position decoding deviation, (c) reconstruction loss, (d) proportion of place fields maintaining original configuration, rotating with perturbation, or remapping. (e) Attention maps for 4 latent variables showing environmental focus. (f-h) Ablation study results: (f) distribution of latent variable paying attention to different environmental features, (g) position decoding deviation, (h) reconstruction loss. Results demonstrate higher importance of proximal cues over distal cues for perspective transformation. Red lines in (b,g,h) indicate random model performance.

First, we evaluated the latent variable position decoding performance. The MLP position decoder was trained on the training set of the original environment and tested on the perturbed environment. Fig. 6b shows the difference in distance between the robot’s predicted and ground truth position under different perturbations for both classification methods. We saw that all three perturbations had significantly greater position deviation than the original in both direction. Moreover, both the arena and the cylinder perturbation performed significantly worse than the background perturbation for the GMP to FPP direction, and the arena perturbation yielded worse performance than the background perturbation for the FPP to GMP direction. This shows once again that the proximal cues on the arena and cylinder were generally more important for the accurate decoding of positions.

Fig. 6c shows the reconstruction loss of each of the three perturbed environments against the original, for both transformation directions. The reconstruction loss due to perturbation of the arena was significantly higher than that of other perturbations for both transformation directions. This demonstrated that the proximal cues from the arena were the most important environmental features for the successful reconstruction of the image for both perspectives. This effect was especially pronounced for the FPP to GMP direction, with the reconstruction loss increased by more than 5X compared to a 2.5X increase for that of the GMP to FPP direction.

Next, we investigated whether there was any remapping of the firing fields for “place cells” in the latent variables of the RNN-VAE. For this test, we compared the place field for the same latent variable before and after the perturbation and used the structural similarity index (SSIM) (36) to find the best match between the perturbed place field and the rotated version of the original place field. SSIM takes a value between −1 to 1, and we used a threshold of 0.2 to determine the best match against counter-clockwise rotation from 0 degrees to 90, 180, and 270 degrees. If none of the SSIM scores for each rotation was greater than 0.2, we deemed this place field as completely remapped and not similar to the original. Fig. 6d shows that for the cylinder perturbation, there was a high proportion of the place cells rotated 90 degrees counter-clockwise, especially in the GMP to FPP direction, matching the same rotation of the environmental feature, which was not found in other perturbations. We note that for the cylinder and arena perturbations there was a significant proportion of place cells that were completely remapped, whereas for the background perturbation, most of the place cells are unchanged. Taken together, this finding shows that proximal cues like the cylinder and arena were more important than distal cues like the background for determining the remapping of place fields. The cylinders were especially important because they could guide the place cells enough that the perturbations caused the place fields to rotate by the same degree.

### Ablation Study

To further investigate what kind of features and cues were important to perspective transformation, we analyzed the results from the Convolutional Block Attention Module (CBAM). Fig. 6e shows selected examples of the attention map of 4 separate latent variables on the same image. The attention map shows which part of the image the latent variable is paying attention to. We can see that some latent variables were paying dominant attention to the background and the floor (left column), while some were paying dominant attention to the cylinder (right column). By analyzing where the dominant attention was deployed to different aspects of the environment by each latent variable, we can interpret the relative importance of each type of environmental cues.

To further analyze the results of the attention mechanism, we aggregated the attention scores for the pixels of the four main types of environmental features, namely the floor, the wall, the cylinder, and the background. This process was automated by looking at the color of the pixels because each of the walls and cylinders had unique colors, and everything else was either background or floor pixels depending on whether they were at the top or bottom half of the image. Fig. 6f shows the distribution of latent variables paying dominant attention to each type of environmental feature within the original and each perturbation. It shows that the majority of the latent variables paid attention to proximal cues such as the floor and the cylinders, followed by the walls, whereas the least attention was given to distal cues such as the background. In addition, we noted that with different perturbations, the percentage of latent variables paying attention to the cylinder increased, especially for the cylinder perturbation. This makes sense because when the cylinders themselves were perturbed, the attention mechanism sought to pay more attention to the cylinders to compensate for the change.

Having established that the latent variables were paying specific attention to each type of environmental feature, we performed an ablation study to investigate the effects of removing latent variables that attend to different environmental features. We set the values of those latent variables to 0 to determine their effect on the overall performance of the model. Fig. 6g shows the same position decoding deviation from the ground truth. It shows that the ablation of latent variables paying dominant attention to “cylinder” introduced the largest increase in position deviation from the original model. This again demonstrated the higher importance of proximal cues such as cylinders on the overall performance of the model. Additionally, Fig. 6h shows the reconstruction loss of the model, with the ablation of the cylinder neurons and the floor latent variables introducing the greatest increase in loss compared to the original. This again demonstrates the importance of the proximal cues to perspective transformation.

## Discussion

In the present work, we suggest that the maintenance of a continuous cognitive map includes switching between perspectives. As we navigate, we are constantly thinking about our location in the world from a visual or other sensory first-person perspective, as well as from a global view of where we are in the world. Unlike prior modeling, which suggests that transformations and linkages could be accomplished through specific encoding of parameters (9, 13, 37), our work is agnostic to what spatial representations support such transformations. Rather, we require a neural network to find a solution to linking FPPs to GMPs and vice versa. Intriguingly, we find many of the same spatial representations (e.g., place cells, head direction cells, border cells, and corner cells) that have been observed in animal studies during two-dimensional maze tasks. Our findings suggest the importance of temporal sequence integration and attention to proximal cues to solve this task. Given the type of spatial responses observed in the model, we suggest that multiple brain regions are contributing to perspective transformations.

### Temporal Sequence is Important for Perspective Transform Modeling

One of the critical insights from our study is the importance of temporal sequence modeling for effective perspective transformation. Unlike static models that rely on individual snapshots of an environment, the RNN architecture in our model allowed it to capture continuous spatial dynamics by incorporating sequences of frames over time. This temporal context significantly improved the model’s performance in transforming between the GMP and the FPP, providing a more biologically plausible mechanism for spatial navigation. By maintaining internal representations across frames, the RNN temporal sequencing model was better equipped to handle the inherent complexities of spatial transformations, including tracking the movement of landmarks and boundaries relative to the agent’s position.

This finding aligns with existing knowledge about how temporal sequences are critical in real-world navigation. In biological systems, animals and humans rely on continuous sensory inputs to track their location as they move through environments, updating their cognitive maps in real time (38, 39). Without the ability to integrate temporal information, momentary spatial disorientation or abrupt perspective shifts could result in significant navigation errors. Our results demonstrate that models lacking this temporal sequence integration, such as static VAEs (22, 23), are less effective at capturing the dynamic nature of perspective transformations, supporting the idea that temporal processing is essential for accurately maintaining spatial coherence over time.

Furthermore, the addition of temporal sequence modeling allowed the latent variables of the model to evolve and adapt as the agent moved through space, mirroring how neural activity in areas like the hippocampus and retrosplenial cortex evolves during navigation (10, 12). This continuous updating of spatial representations is especially important when navigating environments where landmark positions change relative to the agent’s perspective, and when maintaining a coherent sense of direction across frames. Temporal sequence integration, therefore, not only enhances performance in simulated tasks but also reflects a fundamental principle of biological navigation, providing insights that could inform future experimental studies of spatial cognition in both animals and humans.

The importance of temporal information in spatial cognition is further supported by studies on the theta rhythm in the hippocampus, which has been linked to the encoding of sequential spatial information (40). Our RNN-based model captures a similar principle, demonstrating how temporal dependencies can be crucial for accurate spatial representations and transformations. This parallel between our computational model and neurophysiological findings underscores the potential of our approach to shed light on the neural mechanisms underlying spatial navigation and perspective transformation.

### Distributed Cognitive Map of Perspective Transformation

Our model’s latent variables capture many of the representational forms seen in the “distributed cognitive map” of the mammalian brain. This distributed map encompasses not only the hippocampus proper but also the entorhinal cortex, subicular complex, retrosplenial cortex (RSC), posterior parietal cortex, and anterior thalamus (8, 12). Each of these regions contributes unique aspects to spatial cognition, and our model has organically developed analogues to many of these representations.

The emergence of place-cell-like, head-direction-cell-like, and border-cell-like units in our model suggests that these fundamental building blocks of spatial cognition may arise naturally from the computational demands of perspective transformation. This aligns with the concept of parallel map theory, which proposes that spatial representations are built from the integration of multiple neural maps (41). While previous models of spatial navigation have emphasized the role of grid cells in the MEC (39), our results suggest that grid-like representations may not be essential for perspective transformations, particularly in environments with rich landmark information.

Although our model exhibits features reminiscent of hippocampal place cells, it is crucial to note that the hippocampus proper is just one part of the broader network involved in spatial cognition. The combination of border, landmark, and directional tuning we observe is particularly reminiscent of properties seen in the RSC, medial entorhinal cortex (MEC), and subiculum (1, 28, 30). These regions, which integrate multiple forms of spatial information, may be the closest biological analogues to what our model accomplishes.

Our findings suggest that the transformation between global map and first-person perspectives likely demands the full repertoire of tuning properties found in the distributed cognitive map. However, structures that combine border, landmark, and directional information - such as the RSC, MEC, and subiculum - may play a particularly crucial role in this process. The relatively weaker place tuning in our model, compared to the strong place fields typically observed in hippocampal recordings, further supports this interpretation.

Recent studies have highlighted the importance of the RSC in coordinating multiple spatial reference frames (42, 43), a function that aligns closely with the perspective transformation task in our model. The RSC’s role in translating between egocentric and allocentric representations makes it a prime candidate for mediating the types of transformations our model performs.

This work demonstrates the potential for computational models to provide insights into the neural mechanisms underlying complex spatial cognition. By solving a problem that has received relatively little attention in the community - the transformation between global and first-person perspectives - our model offers a new lens through which to view the distributed cognitive map of the brain. Future work could further explore the parallels between our model’s representations and those found in specific brain regions, potentially guiding new hypotheses for neurophysiological investigations.

### Proximal Landmark Cues are More Important than Distal Cue for Spatial Navigation

The introduction of an attention module allowed the model to selectively prioritize specific environmental features, reinforcing the idea that not all spatial information contributes equally to perspective transformation. The attention mechanism revealed a dominant focus on proximal landmarks like the cylinders and walls, which were consistently weighted as important cues for spatial alignment across perspectives. This selective focus, in turn, contributed to the model’s overall performance, as demonstrated by the degradation of performance when these landmarks were rotated or altered during perturbation experiments.

Our results align with neurophysiological studies that have demonstrated the importance of proximal cues in spatial navigation. For instance, O’Keefe and Burgess (44) showed that hippocampal place fields are strongly influenced by the geometric arrangement of nearby boundaries. Similarly, Knierim and Hamilton (45) found that proximal cues exert a more powerful influence than distal cues on place cell firing and spatial behavior.

Rotation perturbation experiments provided additional insight into how the model adapts its internal representations in response to changes in the spatial configuration of landmarks, further highlighting the flexibility and robustness of landmark encoding. This adaptability mirrors the remapping phenomena observed in hippocampal place cells when familiar environments are altered (46).

The results from both the perturbation and ablation experiments strongly indicate that the transformation from GMP to FPP, and vice versa, is critically dependent on these encoded landmarks. Without the proper encoding and attention to these cues, the transformation process becomes less reliable, which is reflected in the increased position decoding error and reconstruction loss. Thus, landmark encoding and the application of an attention mechanism are key computational components of the model, directly contributing to its capacity for successful perspective transformation.

### Implications for Perspective Transformations and Future Human and Animal Experiments

The findings from this study have broad implications for our understanding of spatial cognition and suggest several promising directions for future research in the neuroscience community.

Future studies should further investigate the relative contributions of different brain regions to perspective transformations. Specifically, the roles of the subiculum and RSC in integrating multiple forms of spatial information warrant closer examination by designing experiments that directly compare the involvement of these regions with that of the MEC during GMP-FPP transformations. Furthermore, experiments should look at the temporal evolution of the spatial representations in these brain regions.

To advance our understanding of the neural mechanisms underlying perspective transformations, researchers should consider implementing experiments in both human and animal models. Such studies could involve systematic manipulations of environmental cues, including rotations of local landmarks and global boundaries. This approach would allow for direct comparisons across species and with computational models, potentially revealing conserved mechanisms of spatial cognition.

In human studies, researchers might employ imaging techniques to observe regional differences in brain activity during perspective transformations (47, 48). In animal studies, researchers could utilize high-resolution electrophysiological recordings to examine the remapping of spatial representations at the cellular level (49). By combining these approaches, the field could gain a more comprehensive understanding of how the brain processes and transforms spatial information across different scales and species.

### Conclusion

In conclusion, this study reveals key insights into the computational mechanisms underlying perspective transformations between GMP and FPP. First, incorporating temporal sequence modeling proved crucial, as the recurrent architecture improved the model’s ability to track continuous spatial navigation and perform more accurate transformations by maintaining internal representations across time. Second, our findings challenge the emphasis on grid cells by demonstrating that border and landmark information, along with orientation tuning, plays a more significant role in this perspective shifting task, which likely involves regions such as the RSC and subiculum. Third, the perturbation studies underscored the importance of proximal cues, with nearby landmarks and boundaries having a greater impact on performance than distal cues, suggesting that the brain prioritizes dynamically changing, immediately relevant spatial information in navigation tasks. Finally, these findings provide a computational framework that enhances our understanding of spatial cognition and informs future research in animal and human studies, with a focus on the role of temporal sequences, the distributed nature of spatial representations, and the prioritization of proximal environmental features in perspective transformations.

## Materials and Methods

### Robotic Simulation

The Webot robotic simulation platform (25, 26) is utilized to simulate similar rodents experiments. The Khepera robot is free to explore a 1×1 meter arena with 4 colored walls and 3 colored cylinders. The robot is equipped with 8 proximity sensors for detecting and avoiding walls and obstacles, along with a head-mounted camera capturing 128×128 RGB images. During random exploration, the robot has a 50% chance of moving straight and a 25% chance of veering left or right respectively. Upon detecting a wall or obstacle, there is a 50% chance of either clockwise or counterclockwise turning until the robot’s head direction is away from the obstacle. To expedite the robot’s escape from wall boundaries and prevent prolonged wall-hugging behavior, random targets are placed at the center of the arena, prompting the robot to turn directly towards them.

Pairs of images from the head-mounted camera (FPP) and the top-down view of the simulation are captured every 20 frames, along with the robot’s coordinates, head direction, and motor commands, until 10,000 pairs of images are collected. The simulation environment includes an “entrance hall” background by default, with walls of four separate colors and three cylinders of distinct colors placed in the middle of the arena.

### VAE Training

Variational Autoencoders (VAEs) (23) are constructed to reconstruct images from one perspective to the other, with separate VAEs employed for each direction. All VAEs utilize 50 latent variables. The encoder architecture comprises four layers of convolutional layers with channel sizes of [16, 32, 32, 32] and filter sizes of 3. The decoder architecture consists of five layers of transpose convolutional layers with channel sizes of [512, 256, 128, 64, 32] and filter sizes of 3. Reconstruction loss is measured using standard mean-squared error, and a Kullback-Leibler (KL) divergence term is incorporated to regularize the Gaussian distribution of the latent variables. KL-annealing is implemented to control the relative weight of the KL loss, ranging from 0 to 1, starting from epoch 50 to 1000.

The dataset is divided into 7000 images for training, 1000 images for validation, and 2000 images for testing. Early stopping is enforced when the validation loss does not improve for 100 epochs, following which the best model weights with the lowest validation loss are reinstated. The model demonstrated strong convergence, with the final training reconstruction loss reaching 5.86 × 10^*−*4^ for the FPP to GMP transformation and 4.95 × 10^*−*3^ for the GMP to FPP transformation after 348 epochs of training. The corresponding testing reconstruction losses were 1.78 × 10^*−*3^ and 2.46 × 10^*−*2^ for FPP to GMP and GMP to FPP transformations, respectively. The Adam optimizer with a learning rate of 0.0001 is employed for training. For training the RNN-VAE, only the latent variables are recurrent, while the encoder and decoder remain non-recurrent. The recurrent model is trained using back-propagation through time.

### Cell Type Analysis

To analyze the latent variables for spatial and directional tuning properties, we employed several metrics commonly used in neuroscience literature:

#### Place Cells

Place cell-like activity was quantified using spatial information and spatial coherence as defined by Skaggs et al. (16) and Kubie et al. (15), respectively:

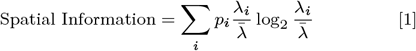

where *p*_*i*_ is the occupancy probability of bin *i, λ*_*i*_ is the mean firing rate in bin *i*, and 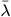 is the overall mean firing rate.

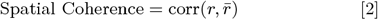

where *r* is the vector of firing rates and 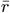 is the vector of smoothed firing rates obtained by averaging over neighboring bins.

#### Head Direction Cells

Head direction tuning was assessed using the Rayleigh vector length as described by Sargolini et al. (31):

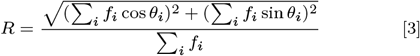

where *f*_*i*_ is the firing rate at heading direction *θ*_*i*_.

#### Border Cells

Border cell activity was quantified using a borderness score following the method of Solstad et al. (30):

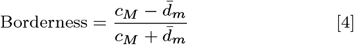

where *c*_*M*_ is the maximum coverage of firing field along any wall, and 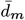 is the mean firing distance to the nearest wall, normalized by half of the shortest side of the environment.

#### Grid Cells

Grid cell-like activity was assessed using a gridness score derived from the spatial autocorrelogram, following the method of Ismakov et al. (32):

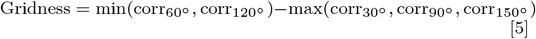

where corr_*θ*_ is the correlation between the autocorrelogram and its rotation by angle *θ*.

#### Object Vector Cells

Object vector cell-like activity was analyzed by computing firing fields as a function of distance and direction to objects in the environment, similar to the approach for border cells, following the methodology established by Høydal et al. (21).

For each cell type, we compared the distribution of scores to established thresholds from rodent literature to classify latent variables as exhibiting properties similar to specific cell types. This analysis allowed us to investigate the emergence of spatially-tuned representations in our model without explicitly incorporating these properties into the architecture.

## Supporting information

Cell Type Ablation Study11

## ACKNOWLEDGMENTS

This work was supported by the Air Force Office of Scientific Research (AFOSR) Contract No. FA9550-19-1-0306, and by the National Institute of Neurological Disorders and Stroke (NINDS) Award No. R01NS135850.

