## Supplementary material for "Neural Mechanisms Linking Global Maps to First-Person Perspectives": Cell Type Ablation Study11

**This PDF file includes:**

Supporting text
Figs. S1 to S2

### Supporting Information Text

### Cell Type Ablation Study

To further investigate the role of different cell types in our model's perspective transformation capabilities, we conducted a cell type ablation study. This study complements our attention-based ablation analysis by focusing on the functional contributions of specific cell types that emerged in our model.

**Methods.** We identified the top 10% of cells belonging to each cell type based on their respective metrics (e.g., spatial information for place cells, head direction score for head direction cells, etc). We then ablated these cells by setting their activity to zero and evaluated the model's performance on two key metrics: position deviation and reconstruction loss.

**Results.** Our ablation study revealed that different cell types contribute differently to the model's performance in terms of reconstruction loss, while position deviation remains relatively stable across cell type ablations. Figures S1 and S2 illustrate the results of our ablation study for both the FPP to GMP and GMP to FPP transformations.

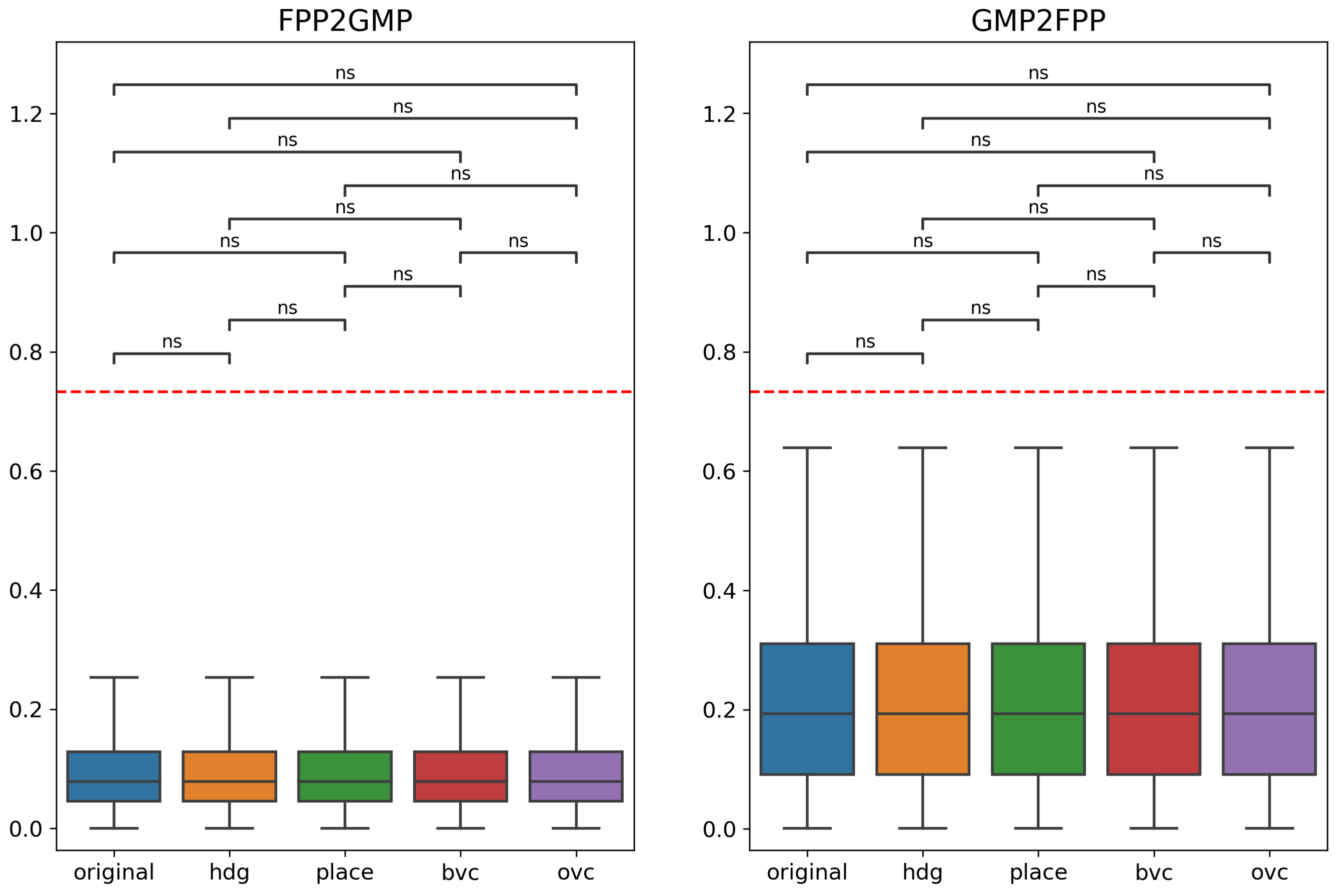


**Fig. S1.** Results of the cell type ablation study showing position deviation. Box plots display the position deviation for the original model and after ablation of the top 10% of cells for each cell type: head direction cells (hdg), place cells, boundary vector cells (bvc), and object vector cells (ovc). Results are shown for both FPP to GMP (left) and GMP to FPP (right) transformations. The red dashed line indicates the performance of an untrained, randomly initialized model. No statistically significant differences were observed between cell type ablations (ns = not significant).

As shown in Figure S1, the ablation of different cell types had minimal impact on position deviation, with no statistically significant differences observed between the original model and any of the ablated versions in either transformation direction. This suggests that the model's ability to maintain accurate position information is robust to the loss of specific cell types.

Figure S2 reveals a more nuanced effect on reconstruction loss. Most notably, the ablation of boundary vector cells (BVC) resulted in a significant increase in reconstruction loss, but only in the FPP to GMP direction. This selective effect suggests that boundary vector cells play a particularly important role in transforming first-person views into map-like representations. Other cell type ablations did not show significant effects on reconstruction loss in either direction.

 Hin Wai Lui, Elizabeth R. Chrastil, Douglas A. Nitz and Jeffrey L. Krichmar

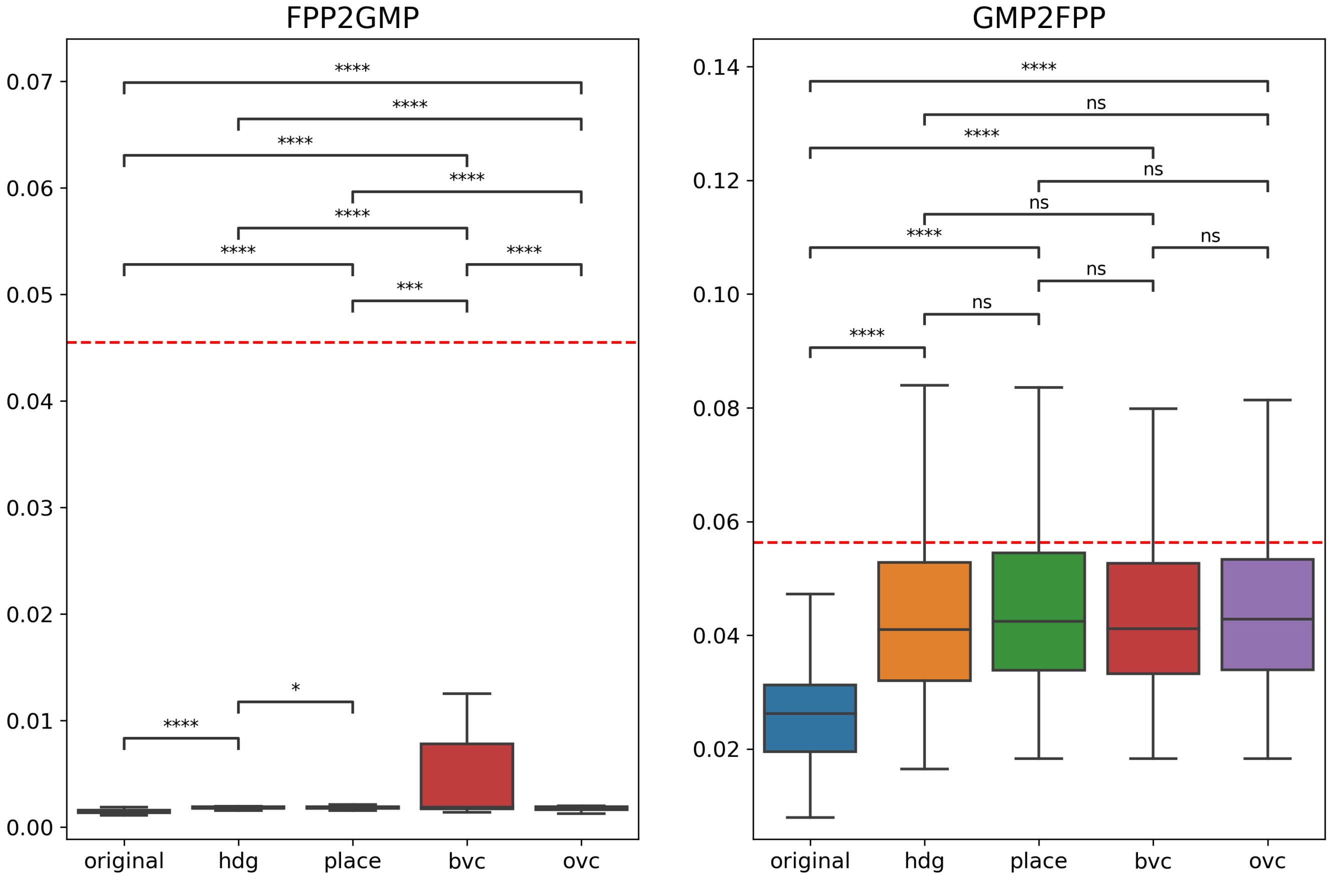


**Fig. S2.** Results of the cell type ablation study showing reconstruction loss. Box plots display the reconstruction loss for the original model and after ablation of the top 10% of cells for each cell type: head direction cells (hdg), place cells, boundary vector cells (bvc), and object vector cells (ovc). Results are shown for both FPP to GMP (left) and GMP to FPP (right) transformations. The red dashed line indicates the performance of an untrained, randomly initialized model. Statistical significance is indicated by asterisks: * $p < 0.05$, ** $p < 0.01$, *** $p < 0.005$, **** $p < 0.001$, ns = not significant. Notably, BVC ablation shows significant effects on reconstruction loss in the FPP to GMP direction.
